# Integrating structural and biological evidence to rerank ESMFold2 protein-protein interactions

**DOI:** 10.64898/2026.09.16.751194

**Authors:** Jindou Xie, Ming Li, Yongping Chai, Guangshuo Ou, Wei Li, Zhengyang Guo

## Abstract

Large-scale protein structure prediction enables proteome-wide protein–protein interaction (PPI) screening, but distinguishing biologically meaningful interactions from spurious interfaces remains challenging. Here, we develop a scalable framework combining fast, MSA-free ESMFold2 prediction with PAE-guided domain parsing and evidence-based reranking. Three-recycle ESMFold2-Fast achieved 57% acceptable-or-better DockQ scores on FoldBench, comparable to AlphaFold2-Multimer while substantially reducing computation. PAE-guided parsing preserved 98.1% of XL-MS cross-links within parsed domain pairs. We then developed the Structure Prediction and Omics informed Classifier (SPOC)-ESMFold2, which integrates structural features with independent biological evidence to prioritize predicted interactions. SPOC-ESMFold2 achieved an AUROC of 0.93 and AUPR of 0.90, compared with 0.87 and 0.79 for a structural-only classifier. Under a 1:128 positive-to-negative ratio, SPOC-ESMFold2 achieved 17.2% recall at 5% false-discovery rate, substantially outperforming structural confidence metrics alone. This framework enables scalable PPI screening by integrating structural plausibility with orthogonal biological evidence to prioritize candidates for experimental investigation.

## Introduction

Recent advances in protein structure prediction have made it possible to generate structural models of protein–protein interactions (PPIs) at a scale that was previously inaccessible to experimental structural biology. AlphaFold2-Multimer and related approaches have enabled systematic prediction of protein complexes, while large-scale applications such as AlphaPulldown and RF-PPI have extended structural interaction prediction toward proteome-wide screens (Bryant et al., 2022; Evans et al., 2022; Yu et al., 2023). These approaches provide a complementary route to experimental methods, and have begun to reveal previously inaccessible protein complexes and interaction mechanisms (Deneke et al., 2024; Deneke et al., 2026).

The emergence of protein language models offers a strategy for scaling structural prediction. ESMFold2 combines the ESMC protein language model with an advanced structure decoder to predict protein structures and multimolecular complexes directly from sequence, without requiring multiple sequence alignments (MSAs) (Candido et al., 2026; Lin et al., 2023). This sequence-based architecture enables efficient GPU inference and makes ESMFold2 particularly attractive for large-scale exploration of protein interaction space. However, increasing the scale of structure generation also exposes a fundamental challenge: how can predicted complexes be distinguished from biologically meaningful interactions when most candidate protein pairs do not interact?

This problem is particularly acute for proteome-scale PPI screening. Fewer than 1% of randomly paired human proteins are expected to interact (Luck et al., 2020)(Luck et al., 2020), creating a highly imbalanced prediction problem in which even a modest false-positive rate can overwhelm true interactions. Moreover, structure prediction methods are intrinsically capable of producing plausible interfaces for non-interacting protein pairs. Consequently, confidence measures derived from the predicted structure—including ipTM and PAE—can identify well-resolved interfaces but do not necessarily establish that the underlying interaction is biologically meaningful. Several approaches have therefore been developed to improve confidence estimation for predicted PPIs, including ipSAE, pDockQ, pDockQ2, local interaction scores (LIS), and measures of prediction consistency (Bryant et al., 2022; Dunbrack, 2025; Kim et al., 2024; Lim et al., 2023; Mirabello & Wallner, 2024).

A recent advance is the Structure Prediction and Omics informed Classifier (SPOC), which addresses this problem by integrating structural confidence with orthogonal biological evidence and evaluating predictions in a recall–false discovery rate (FDR) framework (Schmid & Walter, 2025). This framework is particularly relevant to large-scale interaction discovery because performance is assessed under the severe class imbalance encountered in realistic screening rather than under balanced classification alone. Nevertheless, ESMFold2 differs from AlphaFold2-Multimer in both its sequence representation and structural prediction architecture, and therefore provides a distinct feature space and error profile. Whether the principles developed for AlphaFold-based PPI confidence estimation can be transferred to ESMFold2, and how they should be adapted for efficient proteome-scale screening, remain unclear.

Here, we develop SPOC-ESMFold2, a reranking framework that combines ESMFold2 structural predictions with independent biological evidence to prioritize PPIs under realistic screening conditions. We first establish a computationally efficient ESMFold2-Fast workflow using three recycling iterations and PAE-guided domain parsing, which preserves interaction-relevant structural units while enabling large-scale inference. We then construct a multi-source reference dataset for domain-level PPI classification, integrating experimentally supported interactions from XL-MS and structurally resolved PDB contacts with several complementary negative and decoy sets, followed by stringent structural and homology-based filtering. Using this framework, we show that structural confidence metrics are informative but insufficient under strong class imbalance, whereas integrating structural and biological features substantially improves recall at fixed FDR. SPOC-ESMFold2 therefore provides a scalable approach for prioritizing candidate PPIs from sequence alone, while incorporating orthogonal biological evidence to reduce false discoveries in large-scale interaction screens.

## Results

### Benchmarking ESMFold2-Fast for protein–protein interaction prediction

We first benchmarked ESMFold2-Fast for protein–protein interaction (PPI) prediction using the FoldBench PPI dataset(Xu et al., 2025). We compared a 3-recycle ESMFold2-Fast model with the 10-recycle ESMFold2-Fast and MSA-dependent LocalColabFold, which implements AlphaFold2-Multimer (Evans et al., 2022; Mirdita et al., 2022). For each complex, we generated five predictions and evaluated the model with the highest predicted interface TM-score (ipTM) using DockQ (Figure 1A; Figure S1) (Basu & Wallner, 2016). Despite using single-sequence inputs, 3-recycle ESMFold2-Fast achieved PPI prediction performance comparable to the MSA-dependent LocalColabFold model. Across the 276 complexes analyzed, 57% of predictions reached acceptable-or-better quality (DockQ ≥ 0.23), compared with 60% for 10-recycle ESMFold2-Fast and 56% for LocalColabFold. Notably, the 3-recycle model required substantially less computation, with a total runtime of 266 min compared with 718 min for LocalColabFold. These results indicate that reducing recycling substantially improves computational efficiency while retaining PPI prediction performance, making ESMFold2-Fast suitable for large-scale interaction screening.

**Figure 1.**
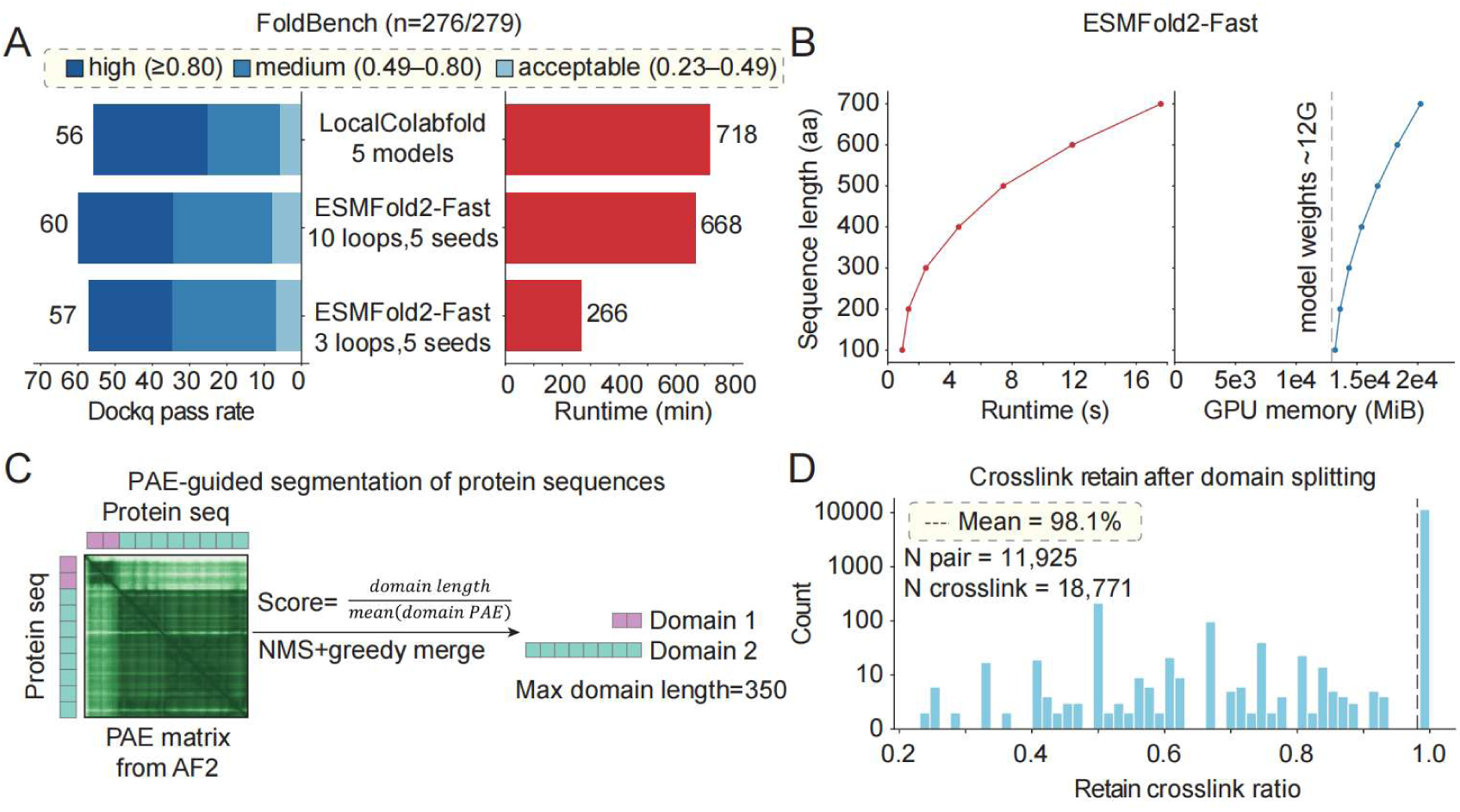
ESMFold2-Fast enables large-scale domain–domain interaction prediction. **(A)** Performance on the FoldBench protein–protein dataset. Horizontal bar plots comparing the computational cost and prediction accuracy of three structure-prediction settings on the FoldBench dataset (n = 276/279): ESMFold2-Fast (3 loops, 5 seeds), ESMFold2-Fast (10 loops, 5 seeds) and LocalColabfold (5 model outputs). DockQ pass rate and runtime (min) are shown in the left and right panels, respectively. In the left panel, bars are colored according to the DockQ score of the predicted structure with the best ipTM across the 5 outputs: high (≥0.80), medium (0.49–0.80) and acceptable (0.23–0.49). **(B)** Left, runtime (s) plotted against sequence length (aa) for ESMFold2-Fast (3 loops). Right, GPU memory usage (MiB) for the same prediction settings; the ∼12G model weights are indicated. **(C)** Schematic of the PAE-guided segmentation strategy. Protein sequences are segmented into structural domains based on the predicted PAE matrix from AFDB, and a segmentation score is calculated for each candidate segmentation. Non-maximum suppression (NMS) and greedy merging are then applied to reach a maximum domain length of 350 aa. **(D)** Distribution of the fraction of crosslinks retained after domain splitting across 11,925 protein pairs and 18,771 crosslinks (mean retained ratio = 98.1%).

### PAE-guided domain parsing preserves interaction-relevant units

We next characterized the computational scaling of ESMFold2-Fast as a function of sequence length. Both inference time and GPU memory usage increased with complex size (Figure 1B). On RTX 4090 GPU with 24 GB VRAM, we therefore imposed a maximum length of 350 residues per chain and 750 residues for the combined sequence. For binary complexes, this corresponds to a maximum combined length of 700 residues when both chains are within the per-chain limit. These constraints defined the sequence-length regime for subsequent large-scale PPI screening and motivated the use of sequence fragments for longer proteins.

Because length-based truncation can disrupt interaction-relevant structural domains, we developed a dynamic domain-parsing algorithm guided by Predicted Aligned Error (PAE) maps from the AlphaFold Database (Varadi et al., 2022) (Figure 1C; Methods). The algorithm identifies flexible inter-domain boundaries based on residue-level PAE and prioritizes candidate domains that are both compact and well resolved. Candidate domains are then selected and merged subject to a maximum length of 350 residues, followed by gap filling to assign the full-length sequence to contiguous domains (Figure 1C; Algorithm 1).

We next asked whether PAE-guided parsing preserves protein regions involved in physical interactions. We benchmarked the resulting domains against cross-linking mass spectrometry (XL-MS) data. For each protein pair, all parsed domain-pair combinations were enumerated, and the pair capturing the largest number of observed cross-links was selected as the representative domain pair. Under this scheme, we found that 98.1% of cross-linking signals remained within a single parsed domain pair across 11,925 protein pairs and 18,771 cross-links (Figure 1D). Thus, PAE-guided parsing largely preserves interaction-relevant sequence units, providing a principled alternative to arbitrary length-based truncation for large-scale PPI prediction.

### A domain-level ESMFold2 PPI dataset from six evidence sources

To establish a high-confidence training and evaluation framework for domain-level PPI prediction, we curated six reference sets from complementary sources, following the reference-set strategy established for SPOC (Figure 2A–F; Methods). Four sets comprised negative or decoy pairs: 80,000 randomly paired human proteins from UniProt (Consortium, 2023), 29,339 non-contact pairs from experimentally resolved PDB complexes (Berman et al., 2000), 1,044 degree-matched random pairs derived from XL-MS datasets, and 10,000 random homodimers (Figure 2A, B, D, E). Two sets comprised positive interactions: unique protein pairs identified from 20 XL-MS studies and contacting pairs derived from PDB complexes deposited after September 30, 2021 (Figure 2C, F), a cutoff chosen to postdate the ESMFold2 training data cutoff (Candido et al., 2026).

**Figure 2.**
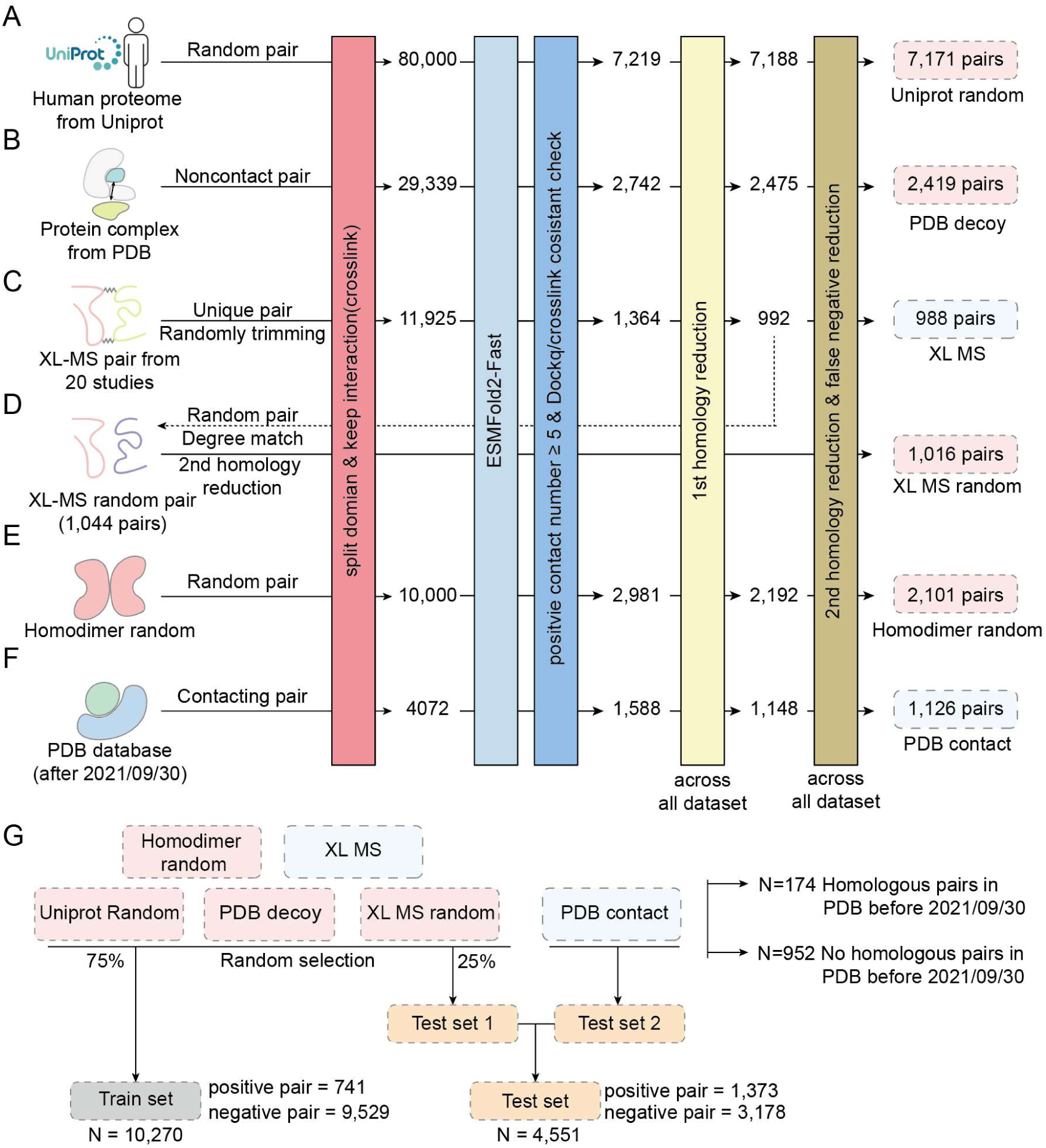
Construction of reference datasets for classifier training and testing. **(A)** UniProt random. 80,000 protein pairs were generated by repeatedly sampling canonical human UniProt entries. **(B)** PDB decoy. Non-contacting pairs, defined as those with fewer than 10 residue pairs having heavy (non-hydrogen) atoms within 5 Å, were extracted from large multi-subunit complexes of known structures in PDB. **(C)** XL-MS. Unique domain–domain cross-link pairs were mined from 20 human or mouse cross-linking studies after random trimming (see Methods). **(D)** XL-MS random. Proteins from the XL-MS set were randomly shuffled and re-paired with matched pairing degrees. After repeated sampling and ESMFold2-Fast folding, >80% of proteins were represented at frequencies within ±1 of those in the XL-MS set. **(E)** Homodimer random. Each protein was paired with itself to generate homodimer decoys. **(F)** PDB contact. Contacting pairs were extracted from protein complexes deposited in the PDB after 30 September 2021, using the complementary criterion of ≥10 residue pairs having heavy (non-hydrogen) atoms within 5 Å. **(G)** Train and test partitions. For each negative dataset (Uniprot random, PDB decoy, XL-MS random, homodimer random), 75% is randomly assigned to the training set and 25% to the test set; for each positive dataset (XL-MS, PDB contact), 25% is randomly assigned to Test set 1. Test sets 1 and 2 are combined into the final test set (1,373 positive and 3,178 negative pairs; N = 4,551); the remaining pairs form the training set (741 positive and 9,529 negative pairs; N = 10,270). For the PDB contact set, N = 174 pairs have homologous matches in the PDB deposited before 2021/09/30, and N = 952 pairs do not.

All candidate pairs were subjected to the same domain-parsing, ESMFold2-Fast prediction, structural filtering, and homology-reduction pipeline (Figure 2). Predicted structures were required to contain at least five positive contacts (Methods). Positive pairs were further filtered for consistency between predicted interfaces and XL-MS cross-links or experimentally determined structures, requiring DockQ > 0.23 where a corresponding PDB structure was available. Finally, two rounds of MMseqs2-based homology reduction (≤30% sequence identity) were applied to minimize training–test leakage, together with removal of negative pairs homologous to any contact pair deposited in PDB (Methods) (Steinegger & Söding, 2017).

After filtering, the six reference sets comprised 7,171 UniProt-random, 2,419 PDB-decoy, 1,016 XL-MS-random, and 2,101 homodimer-random negative pairs, together with 988 XL-MS-supported and 1,126 PDB-contact positive pairs (Figure 2). The two positive sets provide complementary evidence for physical interactions: XL-MS captures interactions in their cellular context, whereas PDB-contact pairs are supported by experimentally resolved structures. We therefore reserved the PDB-contact set exclusively for evaluation, providing an independent positive set that was not used for classifier training.

To further minimize source-specific bias, the classifier was trained using only XL-MS-supported positive pairs. Seventy-five percent of the negative pairs and XL-MS positives were used for training (10,270 pairs; 741 positive and 9,529 negative), while the test set contained the remaining 25% of these pairs together with all 1,126 PDB-contact positives (4,551 pairs; 1,373 positive and 3,178 negative). This design provides both source-independent evaluation and stringent control of homology-mediated information leakage.

### A reranking classifier for ESMFold2 predictions

We next asked whether the quality of ESMFold2-predicted interactions could be improved by integrating multiple structural and biological signals. As a control for potential information leakage, we first trained a graph-based baseline using only network-level properties of the domain–domain interaction graph (Methods). This model performed no better than chance on the held-out test set (AUC = 0.53; AUPR = 0.301), indicating that the graph-based split does not retain substantial network-derived signal across training and test sets (Figure 3A, C; Figure S2). In contrast, the individual structural confidence metrics ipTM, ipSAE, and pDockQ were informative (AUC = 0.85, 0.85, and 0.82, respectively) (Figure 3A, C). However, their performance remained insufficient for reliable discrimination, motivating the integration of additional structural and biological features. We therefore developed a random-forest reranking classifier based on structural features extracted from the predicted interfaces (Breiman, 2001).

**Figure 3.**
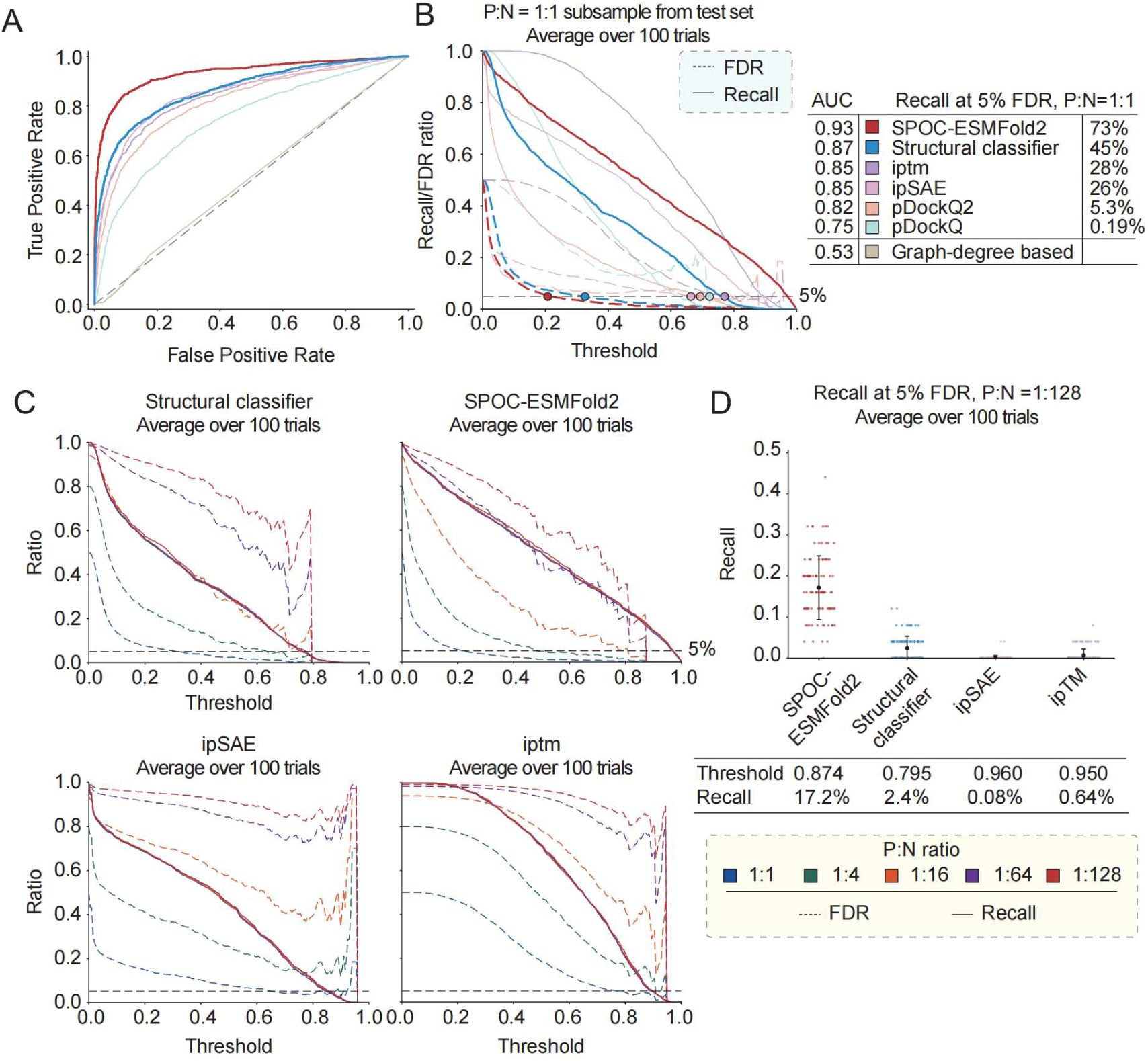
SPOC-ESMFold2 performance on curated test sets. **(A)** ROC curves showing the true positive rate as a function of the false positive rate for SPOC-ESMFold2, the structural classifier, ipSAE and ipTM, averaged over 100 trials. **(B)** Left, recall (solid lines) and FDR (dotted lines) as a function of the score threshold for a P:N = 1:1 subsample of the test set, averaged over 100 trials, for SPOC-ESMFold2, ipSAE, ipTM, pDockQ2, pDockQ, the structural classifier. Right, a table listing the area under the ROC curve (AUC, figure 3A) and the recall at 5% FDR (left panel) for each scoring method. **(C)** Recall and FDR as a function of the score threshold for four scoring methods: SPOC-ESMFold2, the structural classifier, ipSAE and ipTM, at increasing P:N ratios (1:1, 1:4, 1:16, 1:64 and 1:128), averaged over 100 trials. **(D)** Recall at 5% FDR for a P:N = 1:128 subsample of the test set, averaged over 100 trials, for SPOC-ESMFold2, the structural classifier, ipSAE and ipTM; the table below lists the recall at 5% FDR and corresponding threshold for each method.

The structural classifier incorporated 49 features, spanning six categories: ipSAE-derived features, interface features, PAE features, pLDDT features, ipTM features, and other structural features (Table S1). We iteratively pruned uninformative features while retaining complementary structural signals. Unlike the original SPOC framework, we did not include multi-model prediction consistency (avg_models), which requires repeated structure prediction with different sampling seeds and is therefore impractical for proteome-scale screening. Despite this constraint, the resulting classifier integrated information from multiple aspects of the predicted interface rather than relying on a single confidence metric.

We next asked whether orthogonal biological information could further improve interaction ranking. We therefore extended the structural classifier to generate SPOC-ESMFold2 by incorporating genome-scale features independent of the structure prediction, including known interaction evidence from BioGRID, co-expression score from COXPERSdb, DeepLoc 2.0 subcellular localization, CRISPR/BioORCS and DepMap co-essentiality or dependency profiles, T5 protein-language-model embedding similarity, and AlphaMissense pathogenicity scores (Table S2) (Cheng et al., 2023; Elnaggar et al., 2022; Obayashi et al., 2023; Oughtred et al., 2021; Thumuluri et al., 2022; Tsherniak et al., 2017). Feature pruning yielded a final model containing 39 features. SPOC-ESMFold2 thus integrates structural evidence from the predicted complex with independent biological evidence relevant to protein interaction. Rather than relying on a single confidence score, the model provides a unified estimate of interaction likelihood based on the convergence of multiple, partially independent signals.

### The reranking score controls false discovery under screening conditions

We next evaluated classifier performance under the class imbalance expected in proteome-scale screening. Following the recall–FDR framework used in SPOC, we varied the positive-to-negative (P) ratio in the held-out test set from 1:1 to 1:128 and measured recall at a fixed 5% false discovery rate (FDR) (Figure 3B–D; Methods). This range spans balanced evaluation to the highly imbalanced regime expected for single-bait-against-proteome screens, where true interactions are estimated to occur at approximately 1:80. At a balanced 1:1 ratio, SPOC-ESMFold2 recovered 73% of positive pairs at 5% FDR, compared with 45% for the structural classifier, 26–28% for ipSAE and ipTM, and 5.3% for pDockQ (Figure 3B, C). Consistent with these results, SPOC-ESMFold2 achieved an AUC of 0.93 and AUPR of 0.898 on the full test set, compared with 0.87/0.794 for the structural classifier and 0.53/0.301 for the graph-based baseline (Figure 3A, C; Figure S2).

The advantage of integrated reranking became more pronounced under increasing class imbalance. At a N:P ratio of 1:128, SPOC-ESMFold2 retained a mean recall of approximately 0.17 at 5% FDR, compared with approximately 0.02 for the structural classifier and near-zero recall for ipSAE and ipTM (Figure 3C, D). We therefore selected score thresholds of 0.874 for SPOC-ESMFold2 and 0.795 for the structural classifier at the intersection of the 5% FDR criterion and the mean FDR curve across 100 independent 1:128 samplings (Figure 3C, D; Methods). Thus, integrating orthogonal biological evidence substantially improves the ability to control false discoveries when ESMFold2 predictions are evaluated at the extreme class imbalance characteristic of large-scale PPI screening. Hyperparameters for each random-forest classifier were optimized independently by three-fold cross-validation over estimator number, maximum depth, and minimum samples per split, using cross-validated AUPR for model selection (Figure S3; Methods).

### Feature importance reveals complementary structural and biological signals

We next examined the features contributing to classifier performance. In the structural classifier, the most informative individual features were the minimum PAE across interfacial residue pairs (pae_min; Gini importance = 0.067) and the maximum contact score (contact_score_max; 0.058) (Figure 4A), indicating that the classifier directly exploits both interface confidence and the strength of the predicted contacts. The ipSAE feature family collectively represented the largest block of structural importance: although no individual ipSAE metric ranked first, the d0res, d0chn, and d0dom measures and their asymmetry terms each contributed substantially (Gini importance ∼0.02–0.045). Together, these features capture complementary information about the spatial consistency and alignment fidelity of the predicted interface.

**Figure 4.**
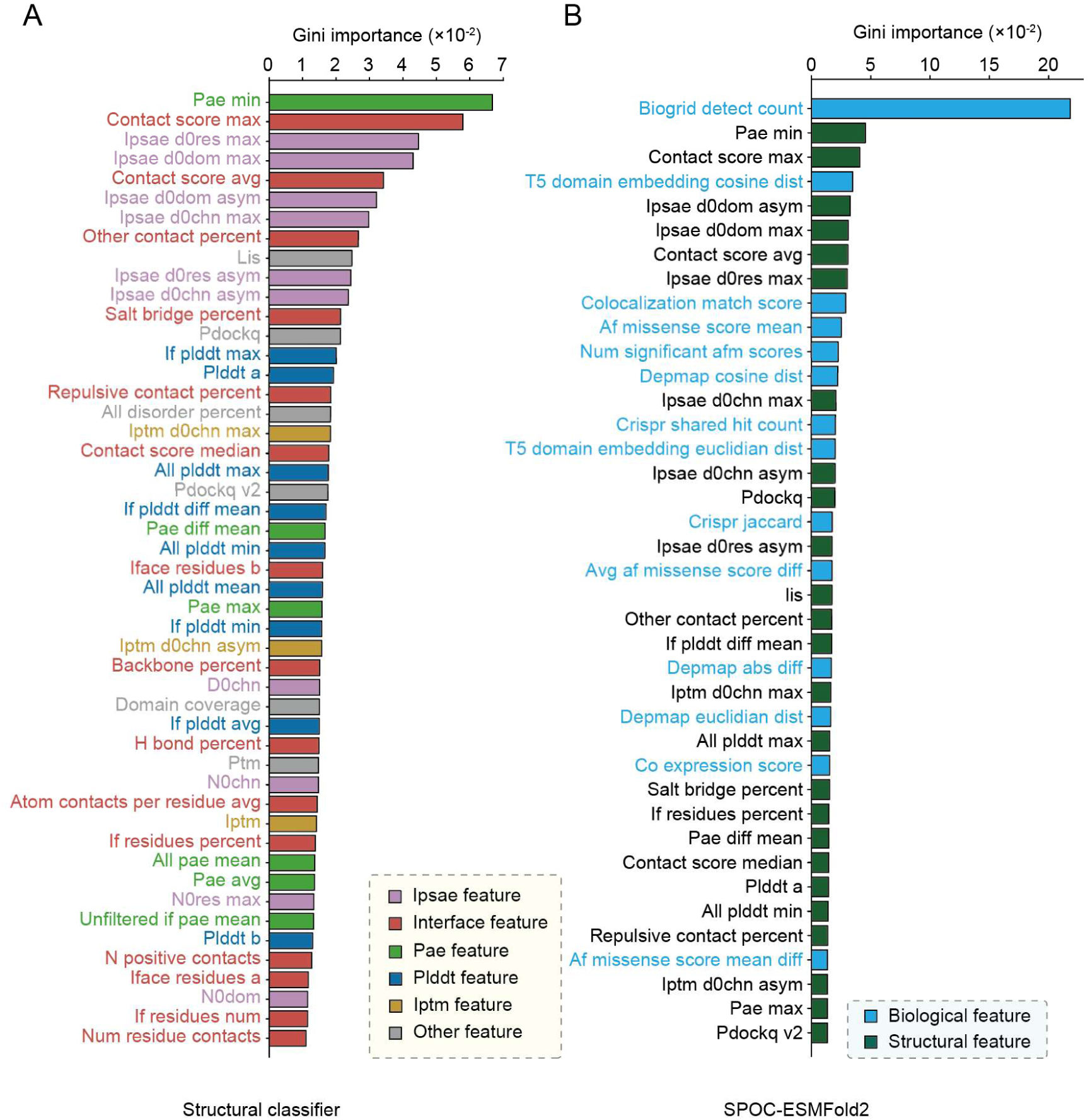
Feature importance of the classifiers. **(A)** Gini importance scores of the features used by the structural classifier. Features are ranked from most to least important and colored by feature category (interface, ipsae, pae, plddt, iptm and other features). **(B)** Gini importance scores of the features used by the SPOC-ESMFold2. Features are ranked from most to least important and colored according to whether they are structural or biological features.

SPOC-ESMFold2 incorporated both structural and genome-scale biological features, with biogrid_detect_count emerging as the most informative individual feature by a substantial margin (Gini importance = 0.21), followed by pae_min, contact_score_max, and T5 domain-embedding similarity (Figure 4B). Overall, structural and biological features accounted for approximately 55% and 45% of total Gini importance, respectively. Thus, the classifier draws on complementary evidence from the predicted interface and independent biological measurements rather than being dominated by either feature class.

## Discussion

The central challenge in large-scale structural interactome mapping is no longer simply generating candidate protein complexes, but determining which predicted interfaces are credible enough to warrant biological investigation. Structure prediction and interaction prediction are related but distinct problems: a protein pair can adopt a geometrically plausible interface in silico without forming an interaction in the cell. This distinction becomes critical as screening scales increase, because even a small false-positive rate can overwhelm genuine interactions when most protein pairs are non-interacting. Our results show that this problem cannot be adequately addressed by a single measure of structural confidence. Instead, reliable prioritization requires combining information about the predicted interface with evidence that is independent of the folding model.

This observation places SPOC-ESMFold2 in a broader progression from structure prediction toward evidence-based interaction inference. Previous methods have demonstrated that AlphaFold-based models can be used to screen large numbers of protein pairs, while SPOC established that calibrated integration of structural and biological features can improve discrimination under realistic screening conditions (Bryant et al., 2022; Schmid & Walter, 2025). Our results extend this principle to ESMFold2, despite its substantially different prediction architecture and its reliance on protein-language-model representations rather than MSAs. Importantly, the transfer is not achieved by applying AlphaFold-derived thresholds to ESMFold2. Instead, the structural features must be evaluated and calibrated against the characteristic outputs of the new predictor. This suggests that the distinction between structure confidence and interaction confidence is general, whereas the quantitative features used to estimate it are predictor-specific.

The computational efficiency of ESMFold2-Fast further changes the practical regime in which structural PPI prediction can be performed. With three recycling iterations, its FoldBench performance was comparable to that of LocalColabFold AlphaFold2-Multimer while requiring substantially less computation. This advantage is particularly relevant for interaction screening, where the objective is not necessarily to obtain the most refined structure for every candidate pair, but to evaluate very large numbers of candidate interfaces sufficiently well to enable effective prioritization. In this setting, a modest reduction in per-complex accuracy may be outweighed by the ability to screen a substantially larger search space. The results therefore support a division of labor in which fast structure prediction generates candidate interaction hypotheses, while downstream reranking determines which hypotheses merit further attention.

The need for domain-level modeling is a related consequence of scaling structure prediction to the proteome. Fixed-length truncation is computationally convenient but can arbitrarily divide interaction-relevant regions. Our PAE-guided parsing provides a data-driven alternative by using the predicted structural organization of each protein to define compact sequence units. The retention of 98.1% of XL-MS cross-links within parsed domain pairs indicates that these boundaries largely preserve experimentally supported interaction units. This result is encouraging because it suggests that sequence-length constraints need not be treated simply as a limitation of the prediction engine; they can instead be incorporated into a biologically informed decomposition of the proteome. Nevertheless, XL-MS provides evidence for physical proximity rather than a complete definition of functional domains, and the performance of the parsing strategy should ultimately be evaluated against additional structural and functional annotations.

Our classifier results further clarify why structural confidence alone has limited resolving power. Metrics such as ipTM, ipSAE, and pDockQ are informative because they quantify properties of the predicted interface, but they remain fundamentally agnostic to much of the biological context in which interactions occur. The feature-importance analysis reinforces this point: ipSAE and direct interface-confidence measures constitute substantial components of the structural signal, yet adding orthogonal biological information produced a marked improvement in ranking, particularly under extreme class imbalance. Thus, the value of SPOC-ESMFold2 is not simply that it contains more features. Rather, it combines different types of evidence that constrain different aspects of the interaction hypothesis. Structural features ask whether an interface is plausible; biological features provide independent evidence that the two proteins occupy compatible cellular or functional contexts.

The prominence of BioGRID evidence illustrates both the strength and the caveat of this approach. Prior interaction evidence is highly informative, as expected, and its contribution indicates that existing biological knowledge can substantially improve the prioritization of structural predictions. At the same time, this means that the highest-confidence predictions are not necessarily the most novel ones. A classifier trained on known interactions will naturally favor candidates that resemble interactions already supported by experimental evidence. Consequently, the principal value of SPOC-ESMFold2 for discovery should not be judged solely by its ability to recover known interactions, but also by whether its ranking can enrich experimentally validated interactions among previously uncharacterized pairs. This distinction will be important when assessing the method’s capacity for genuinely de novo interactome discovery.

Our evaluation framework was designed to address this issue as far as possible. By training on XL-MS-supported positives while reserving the post-cutoff PDB-contact set for testing, we evaluated generalization to a positive evidence source that was not used for training. The recall–FDR analysis is also important because conventional AUROC values can obscure the practical difficulty of highly imbalanced interaction screening. The approximately 1:128 evaluation regime represents a substantially more demanding setting than balanced classification, and the persistence of recall at 5% FDR demonstrates that the integrated score retains useful discriminative power when false positives greatly outnumber true interactions. We therefore consider low-FDR performance under realistic class imbalance a more informative measure of utility for large-scale PPI screening than balanced accuracy alone.

Several limitations define the current scope of the framework. First, we deliberately omitted multi-model prediction consistency, an informative feature in the original SPOC framework, because obtaining it would require repeated ESMFold2 predictions and substantially increase computational cost. This choice favors throughput but leaves potentially useful information about prediction reproducibility unexplored. An important future direction will be to determine whether a limited number of additional samples can provide sufficient consistency information to improve ranking without sacrificing the scalability of the pipeline. Second, our current implementation is constrained by sequence length and was designed primarily for binary human protein interactions. Extending the approach to larger multidomain proteins, membrane-associated complexes, and higher-order assemblies will require further development of domain decomposition and interface scoring. Third, the reference sets necessarily provide an imperfect approximation to the true interaction landscape. In particular, the negative sets represent different forms of decoy or non-contact pairs rather than a definitive catalogue of biological non-interactions. Experimental validation will therefore remain essential for estimating the absolute precision of predictions, especially among previously unsupported candidates.

More broadly, the framework suggests a useful strategy for future computational interactomics. Rather than treating a structure prediction as an endpoint, structural models can be viewed as hypotheses that can be evaluated against independent layers of biological evidence. This perspective is particularly powerful as protein-language-model-based predictors make structure generation increasingly fast and scalable. The limiting factor may consequently shift from the generation of structural models to the principled prioritization of the enormous number of candidate interactions that can now be generated. Integrating structural, genetic, expression, localization, evolutionary, and other functional evidence provides one route toward addressing this problem while maintaining an explicit estimate of false-discovery risk.

The present study therefore establishes a framework in which fast MSA-free structure prediction and evidence-based statistical prioritization are complementary rather than competing approaches. ESMFold2-Fast expands the feasible scale of structural hypothesis generation, PAE-guided parsing preserves interaction-relevant sequence units, and SPOC-ESMFold2 uses orthogonal evidence to distinguish more credible interaction hypotheses from plausible but unsupported interfaces. The next test of this framework is experimental: whether its low-FDR ranking can enrich genuinely novel interactions in large-scale validation experiments. If so, this combination of scalable structure prediction and calibrated evidence integration could provide a practical route toward more systematic mapping of the human interactome.

## Acknowledgments

This work was supported by the National Key R&D Program of China and the National Natural Science Foundation of China.

## Data and code availability

The full ESMFold2 model outputs, reference datasets, feature matrices and reproducibility scripts are available on https://github.com/Tttgb/SPOC_ESMFold2.

## Author Contributions

J.X., G. O. and Z. G. designed research; J.X., M. L., Y.C., G. O. and Z. G. performed research; J.X., M. L., Y.C. and Z. G. analyzed data; and J.X., M. L., G. O., W.L. and Z. G. wrote the paper.

## Supplementary Materials

### Methods

Supplementary Table S1, 49 features computed from the predicted interface used by structural classifier.

Supplementary Table S2, 39 features (25 structural + 14 biological) used by SPOC-ESMFold2.

Supplementary Figure S1, Per-output DockQ performance on FoldBench.

Supplementary Figure S2, Precision–recall performance of the reranking classifiers.

Supplementary Figure S3, Hyperparameter tuning of the random-forest classifiers by cross-validated grid search.

Supplementary Figure S4, The definition of positive contacting residue.

### Methods

#### ESMFold2 inference

All pairs were folded with ESMFold2-Fast (ESMC 6B encoder, 3 recycling loops, 50 sampling steps) on 8× RTX 4090 GPUs, constraining complexes to ≤750 AAs and single chains to ≤350 AAs per GPU memory. Per-pair outputs were the predicted coordinates (PDB/MMCIF), per-residue pLDDT (0–1), the PAE matrix (Å, capped ∼31.7), and global iptm/ptm.

#### Domain parsing

To define interaction units at the domain level, we first segmented each protein into compact structural domains using its predicted aligned error (PAE) matrix (Figure 1C). After symmetrizing the matrix, we computed a residue-level boundary score: for every position i we averaged the PAE over the cross-block between the two flanking 30-residue windows (i−30…i−1 vs. i…i+30), and local maxima of this profile (minimum prominence 2 Å, ≥20 residues apart) were taken as candidate cut points, since high inter-window PAE indicates low confidence in the relative orientation of the two halves and thus marks an inter-domain hinge. Candidate domains were enumerated as all intervals of 80–350 residues between any pair of cut points, together with all intervals between adjacent cut points (≤350 residues), and were ranked by a compactness score equal to the domain length divided by its mean intra-domain PAE (+ε = 10⁻⁵ for numerical stability), which favors long segments whose residues are consistently well aligned internally. If no candidate satisfied the length constraints, the sequence was instead tiled into 350-residue windows with 100-residue overlap. To obtain a final, non-redundant segmentation, we applied one-dimensional non-maximum suppression, in which candidates whose intersection-over-minimum-length with an already-retained, higher-scoring domain exceeded 0.5 were discarded. Surviving domains were then merged by a left-to-right scan in which the running span was extended as long as its total length did not exceed 350 residues. Uncovered stretches were patched by windows extending 100 residues beyond each gap on both sides; gaps longer than 350 residues were tiled into ≤350-residue chunks overlapping by 100 residues, so that every residue is assigned to a contiguous, compact structural domain with 100% sequence coverage. Proteins no longer than 350 residues were kept intact as a single domain without further segmentation (Algorithm 1).

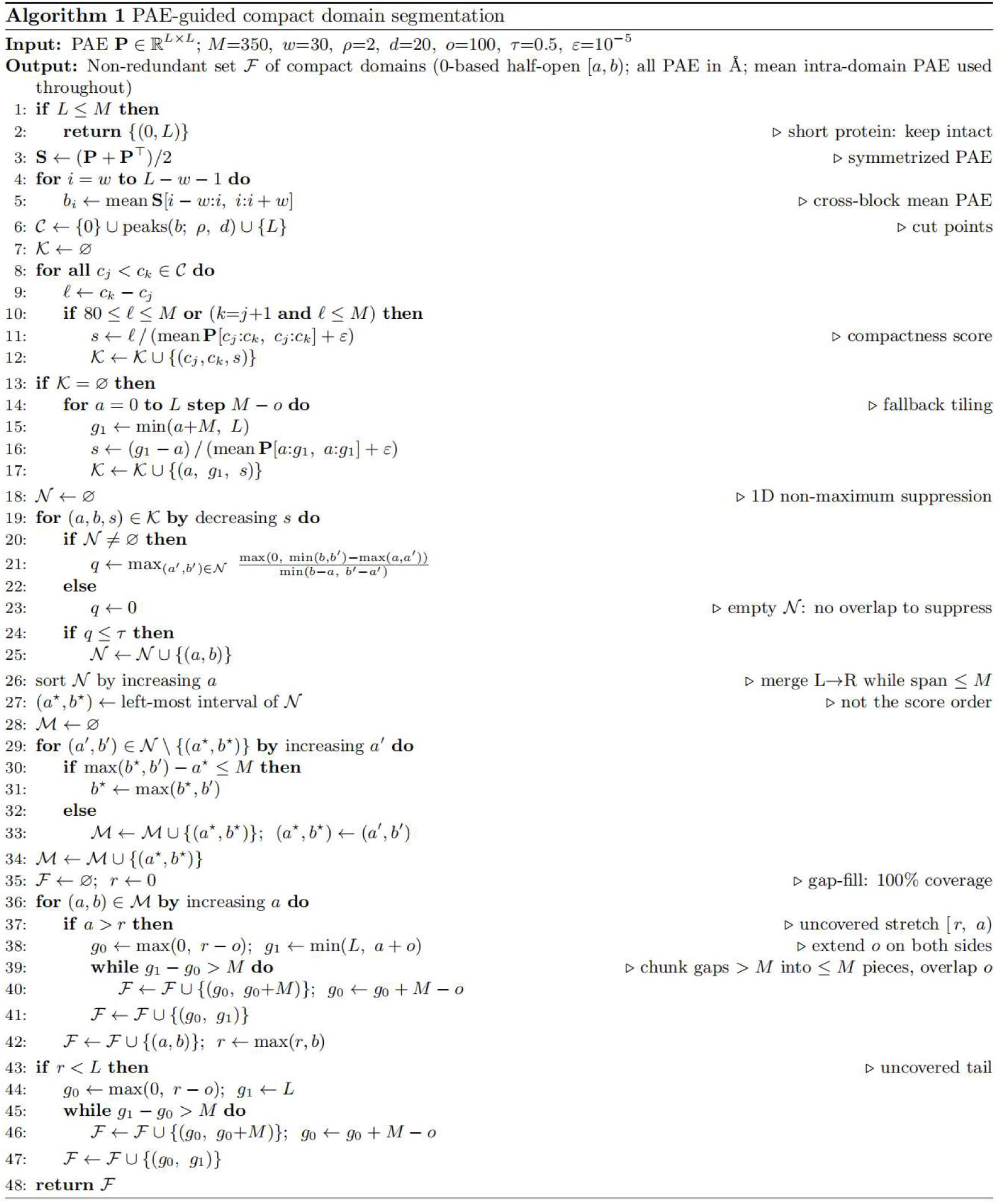

### Uniprot random reference set building

For the random human-protein-pair set, negatives were sampled from the human proteome: the full-length sequences of the human reviewed proteome (UniProtKB/Swiss-Prot) were repeatedly shuffled and paired into distinct protein pairs (self-pairs excluded), generating 100,000 candidate protein pairs (Figure 2A). Because such pairs carry no cross-link or contact evidence with which to guide domain selection, each protein was segmented into structural domains (Methods) and a single domain–domain pair was drawn uniformly at random from all possible domain pairs of the two proteins. Domain splitting requires an AlphaFold-model PAE matrix, and proteins >2,600 aa lacking an AlphaFold structure cannot be split; because such proteins are removed during segmentation, we passed all 100,000 candidate pairs through domain splitting and then drew 80,000 domain–domain pairs uniformly at random from those successfully generated, fixing the number of random negatives modelled with ESMFold2 at 80,000 (Figure 2A). After contact-positivity filtering and homology reduction, 7,171 random human protein pairs were retained.

### PDB decoy reference set building

For the PDB-decoy set, we used pairs of human proteins that co-occur within the same PDB complex but do not physically interact. For every PDB entry containing at least three human chains, all pairs of distinct human proteins present in that entry were recorded as co-occurring; two chains were considered contacting under the same criterion as for the PDB-contact set (≥10 contacting residue pairs with heavy atoms < 5 Å). A co-occurring protein pair was classified as non-interacting only if it was never observed to contact in any PDB entry, i.e., the two proteins are captured in the same complex yet never form a direct interface (31,557 candidate pairs). Because decoy pairs carry no defined interface, each protein was segmented into structural domains and each candidate was represented by a single domain–domain pair drawn uniformly at random from all possible domain pairs of the two proteins, as for the random set (Figure 2B). After ESMFold2-Fast modeling, contact-positivity filtering and homology reduction, 2,419 non-interacting chain pairs were retained.

### XL-MS reference set building

For the XL-MS set, inter-protein cross-links were compiled from 20 published cross-linking–mass spectrometry (XL-MS) studies (same as SPOC) (Schmid & Walter, 2025) and assigned to full-length protein sequences. Peptide–protein assignments originating from non-human organisms were transferred to their human orthologues when a confident mapping could be made, and any pair that could not be represented on human sequences was discarded. Cross-links were then classified as intra-or inter-molecular from the identities of the two cross-linked proteins and the two cross-linked residue positions. When a cross-link joined two different proteins, it was kept as hetero-oligomeric evidence of an interaction. When the same UniProt protein appeared on both sides of a cross-link, we used the cross-linked residue positions to distinguish a self-link from a homodimer link: if the residue on one side was cross-linked to the same residue on the other side (i.e., a residue would be cross-linked to itself), the link cannot be formed within a single chain and was therefore assigned to two copies of the same protein, being kept as a homodimer (“dimer”) link; if the two sides mapped to different residues, the link was treated as an intra-molecular self-link (“self”) within a single chain and was removed, as it does not report on protein–protein interactions. The remaining inter-molecular cross-links (both homo-and hetero-oligomeric) were collapsed to the protein-pair level, giving 21,277 unique cross-linked protein pairs. Because a small number of extensively studied proteins dominate XL-MS compendia, we trimmed over-represented proteins before domain splitting: iteratively, the most-common protein across all pairs was identified and, of all pairs containing it, all but 28 randomly chosen pairs were removed; this was repeated until no protein occurred in more than 28 pairs (21,277 → 13,908 pairs). Histone proteins, identified using a list of human histone identifiers, were then removed at random until histone-containing pairs represented only 1% of the set (→ 13,375 pairs); Each remaining pair was represented at the domain level by segmenting both full-length sequences into structural domains and choosing the domain–domain pair that contained the most cross-linked residue pairs, i.e., that captured both residues of the largest number of cross-links (Figure 2C); protein pairs for which no domain pair captured a cross-link were discarded. The representative domain pairs were modeled as binary complexes with ESMFold2-Fast and retained only if the model was contact-positive and recapitulated the cross-link, i.e., contained at least one cross-linked residue pair whose Cα atom lie within 36 Å of each other in the model (crosslink consistence check), yielding 988 XL-MS positives (Figure 2C).

### XL-MS random reference set building

For the XL-MS random set, we designed negatives that mirror the composition of the XL-MS positives after 1st homology reduction, in which a minority of proteins are heavily represented. For every protein in the XL-MS set we counted its degree, i.e., the number of pairs in which it participates. 45,000 candidate negatives were generated directly at the domain level: two domains were drawn at random from the domain pool of the XL-MS positives after 1st homology reduction, with sampling weighted by the degree of their source protein so that the expected number of pairs per protein reproduced its XL-MS degree; self and duplicate domain pairs were excluded (Figure 2D). Because candidates were drawn directly as domain–domain pairs, each sampled pair served as one negative without further domain splitting. Candidates were modeled with ESMFold2-Fast and required to be contact-positive, and a greedy procedure selected a subset in which >80% of the XL-MS proteins were matched. Then, after 2nd homology reduction, 1,016 negatives were generated (a protein being considered matched only if it occurred at least once in the negative set and its occurrence count differed from its XL-MS count by at most ±1) (Figure 2D).

### Homodimer random reference set building

For the random-homodimer set, proteins were sampled at random from the human reviewed proteome and each sampled protein was combined with itself to form a homodimer pair. Each protein was segmented into structural domains and a single domain was selected at random; the two chains of the homodimer were then built from two identical copies of that one domain, i.e., domain selection was one-to-one (one randomly chosen domain per protein, paired with itself) rather than sampling from all possible pairwise combinations of the protein’s domains. Because domain splitting requires an AlphaFold-model PAE matrix and proteins >2,600 aa lacking an AlphaFold structure cannot be split, we oversampled 12,000 proteins and, after discarding proteins that could not be segmented, retained 10,000 homodimer domain pairs, fixing the number of random homodimers modelled with ESMFold2 at 10,000. Pairs exceeding the sequence-length limit for modeling were discarded. After ESMFold2 modeling, contact-positivity filtering and homology reduction, 2,101 random homodimers were retained.

### PDB contact reference set building

For the PDB-contact set, experimentally determined heteromeric interactions were derived from the Protein Data Bank (PDB). We considered PDB entries containing human protein chains whose initial deposition date was after 30 September 2021 and resolution < 3.5 Å, thereby excluding complexes that may overlap with ESMFold2-Fast training data; human chains were mapped to UniProtKB/Swiss-Prot sequences. Two chains were defined as contacting when they shared ≥10 residue pairs containing at least one pair of heavy atoms < 5 Å apart, and we retained heteromeric chain pairs corresponding to two different proteins (Only contacting pairs both from homo sapiens are considered). When a protein pair occurred in several PDB entries, the entry with the largest number of contacts was used as the representative structure, and its interacting residues were mapped onto the full-length UniProt sequences. Then, each sequence was then segmented into structural domains, and the domain–domain pair containing the most contacting residue pairs was selected as the representative (Figure 2F). The representative domain pairs were modeled with ESMFold2 and retained only if the model was contact-positive and superimposed well onto the native complex (DockQ score > 0.23), giving 1,126 positive PDB heterodimers (Figure 2F).

### Contact-positivity

A predicted structure was contact-positive if it had ≥5 interfacial residue pairs with PAE < 15 Å, both residues with pLDDT > 0.5, and inter-residue separation 1–5 Å (Figure S4).

### Dockq check

DockQ scores were calculated with the DockQ Python package (Basu & Wallner, 2016), and a model was retained only if DockQ > 0.23, indicating that the predicted domain–domain complex superimposed acceptably well onto the native structure (the corresponding PDB entry trimmed to the two interacting chains).

### 1st homology reduction

After contact-positivity filtering, sequence redundancy was removed across the five sets generated up to this point — random human-protein-pair, PDB-decoy, XL-MS, PDB-contact and random-homodimer — by clustering all domain sequences with MMseqs2 at ≥30% sequence identity and ≥50% coverage (of both sequences). A pair was discarded when its two domains fell into the same sequence clusters as the two domains of an already retained pair, i.e., when it shared ≥30% sequence identity with both partners of another pair. When homologous pairs carried different labels, positive pairs were preferentially retained over negative ones.

### 2nd homology reduction, false negative reduction

After the XL-MS random set is generated from XL-MS set after 1st homology reduction, the same MMseqs2 homology reduction was repeated across all six sets. In addition, false negatives were removed from the negative sets: we compiled a reference of interacting chains from all PDB structures containing human chains, identifying every pair of chains that contact in a deposited structure (human chains mapped to UniProt sequences; other chains identified by their sequence). Each domain of every negative pair was then searched against this reference with MMseqs2 (≥30% sequence identity, ≥50% coverage). A negative whose two domains each matched chains that form a PDB-interacting pair — i.e., a pair with >30% similarity to a known PDB interaction — was flagged as a likely true interaction that had been mislabeled as negative and was removed.

### Train/test split

75% of each negative set and 75% of XL-MS formed the training set; the remaining 25% of negatives together with all PDB contacts formed the test set (Figure 2G).

### Feature extraction

Per-pair features were computed from the predicted structure and its confidence outputs. Structural features (49; Table S1) fall into six categories: interface, pLDDT, PAE, ipTM, ipSAE and other features. Interface features characterize the geometry and chemical composition of the predicted interface: the number of positive-contact residue pairs and of unique inter-residue contacts, interfacial residue counts and fractions per chain and in total, the mean number of atom contacts per interfacial residue, per-pair contact scores that weight the number of atom contacts by residue confidence, and chemical descriptors of the interfacial contacts. For the chemical descriptors, inter-chain heavy-atom pairs within 5 Å were classified as backbone contacts (involving a main-chain N, CA, C or O atom), hydrogen bonds (N-O pairs within 3 Å), salt bridges (between residues of opposite net charge), repulsive contacts (between residues of like net charge) or other contacts, with contacts closer than 1 Å treated as steric clashes and their residues excluded from the downstream statistics; their fractions of all interfacial atom contacts were recorded (backbone_percent, h_bond_percent, salt_bridge_percent, repulsive_contact_percent and other_contact_percent). pLDDT features comprise chain-level and interface-level statistics of the predicted per-residue confidence (all_plddt_mean/max/min, plddt_A/B and if_plddt_avg/max/min/diff). PAE features summarize the predicted aligned error over all residue pairs and over interfacial residue pairs, including the mean/min/max and the asymmetry between the two PAE directions (all_pae_mean, unfiltered_if_pae_mean, pae_avg/max/min and pae_diff_mean). ipTM features comprise ipTM and the ipTM-like d0chn-scaled scores (iptm, iptm_d0chn_asym, iptm_d0chn_max). The ipSAE features comprise the PAE-based interface-confidence scores ipSAE with per-residue, per-domain and per-chain d0 scaling together with the associated d0/n0 scales (Dunbrack, 2025). The remaining structural features include pDockQ (v1/v2), pTM, domain coverage, LIS and the fraction of disordered residues (pLDDT < 50) (Bryant et al., 2022; Mirabello & Wallner, 2024).

The contact score of an interfacial residue pair x–y is the number of atom contacts between the two residues (heavy atoms < 5 Å apart) multiplied by their averaged pLDDT and divided by 1 plus the average of the two directional PAE values, PAE(x, y) and PAE(y, x) (Schmid & Walter, 2025):

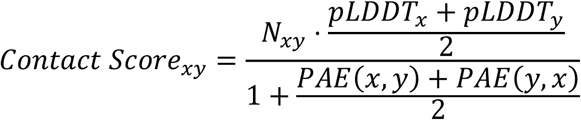

Biological features (n=14; Table S2) combine interaction and functional evidence for the two parent proteins: BioGRID interaction evidence count (https://downloads.thebiogrid.org/) (Oughtred et al., 2021), Coexpression (https://coxpresdb.jp/) (Obayashi et al., 2023), DeepLoc 2.0 co-localization (Thumuluri et al., 2022), BioGRID CRISPR (ORCS) screens and Jaccard similarity (https://downloads.thebiogrid.org/) (Oughtred et al., 2021), DepMap genetic-dependence distances (https://depmap.org/portal/home/) (Oughtred et al., 2021), sequence-embedding distances between the two domains from a protein language model (ProtT5) (https://www.uniprot.org/help/downloads) (Elnaggar et al., 2022) and AlphaMissense pathogenicity statistics (https://alphamissense.hegelab.org/download) (Cheng et al., 2023) over the interface.

### Random forest training

Random forests were fitted separately to the 54 structural features and to the combined set of 68 structural and biological features, after a fixed stratified train/test split (labels stratified, random_state 42): 75% of the random, PDB-decoy, XL-MS, XL-MS-random and random-homodimer pairs formed the training set (10,270 pairs) and the remaining 25%, together with the entire PDB-contact set, formed the held-out test set (4,551 pairs), which was used only for the final evaluation. Missing values were imputed with the per-feature median, with the imputer fitted on the training set alone. Hyperparameters were selected by three-fold stratified cross-validation (shuffled, random_state 42) optimizing average precision (sklearn average_precision_score) over n_estimators ∈ {200, 400, 600, 1000}, min_samples_split ∈ {2, 5, 10, 20}, bootstrap ∈ {true, false} and criterion ∈ {gini, log_loss}, and over max_depth ∈ {8, 12, 16, 20, 24} for the integrated model and {5, 8, 12, 16, 20} for the structural model (Figure S4).

Features were then pruned iteratively: in each round the best cross-validated model was refitted on the full training set, features whose Gini importance was < 0.01 were removed, and model selection was repeated on the remaining features until no feature was pruned. This retained 49 structural features for the structural classifier (Table S1) and 39 features (25 structural + 14 biological) for the integrated classifier (Table S2). The final models were refitted on the full training set with the selected hyperparameters (structural: 400 trees, max depth 12, min_samples_split 10; integrated: 600 trees, max depth 12, min_samples_split 10; both bootstrap = false, criterion = log_loss, unweighted classes, default max_features). Test AUPR was 0.794 for the structural and 0.898 for the integrated classifier.

Random forests were fitted separately to the structural features and to the combined structural and biological features. The structural classifier contained 49 features, and the integrated classifier retained 39 after iterative pruning using a Gini-importance threshold of 0.01. Hyperparameters were selected by three-fold cross-validated AUPR over n_estimators ∈ {200, 400, 600, 1000}, max_depth ∈ {8, 12, 16, 20, 24}, and min_samples_split ∈ {2, 5, 10, 20} (Figure S4). The selected integrated-model configuration was 600 estimators, depth 12 and a minimum of 10 samples per split; the structural model used 400 estimators, depth 12 and a minimum of 10 samples per split. Missing feature values were imputed.

### Graph-structure baseline model

To examine whether the evaluation of our structure-based model is inflated by *protein overlap* between training and test sets, we construct a baseline that relies solely on interaction-network topology. Two undirected unweighted graphs are built from the training set: *G*^+^, where an edge connects two proteins labeled as interacting (label = 1), and *G*^all^, where an edge connects any co-occurring protein pair. For each vertex we compute its degree deg(*v*), i.e., the number of distinct interaction partners. For a protein pair (*A*, *B*), four basic degree features 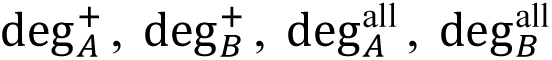 are extracted and combined element-wise via sum, absolute difference, min, max and product, yielding 14 topological features. Then, A random forest is trained on these features with AUPR as the tuning objective.

### Recall–FDR analysis

Recall–FDR analysis follows SPOC. For each P:N ratio from 1:1 to 1:128, the class imbalance was adjusted in two regimes: at moderate ratios, negatives were subsampled with replacement to N = ratio × P; at extreme ratios, positives were subsampled while holding the full negative set (n = 3,178) fixed, as the negative pool is too small to support high replication factors. Recall was reported at a 5% FDR, and results were averaged over 100 resampling trials.

---

**Figure S1.**
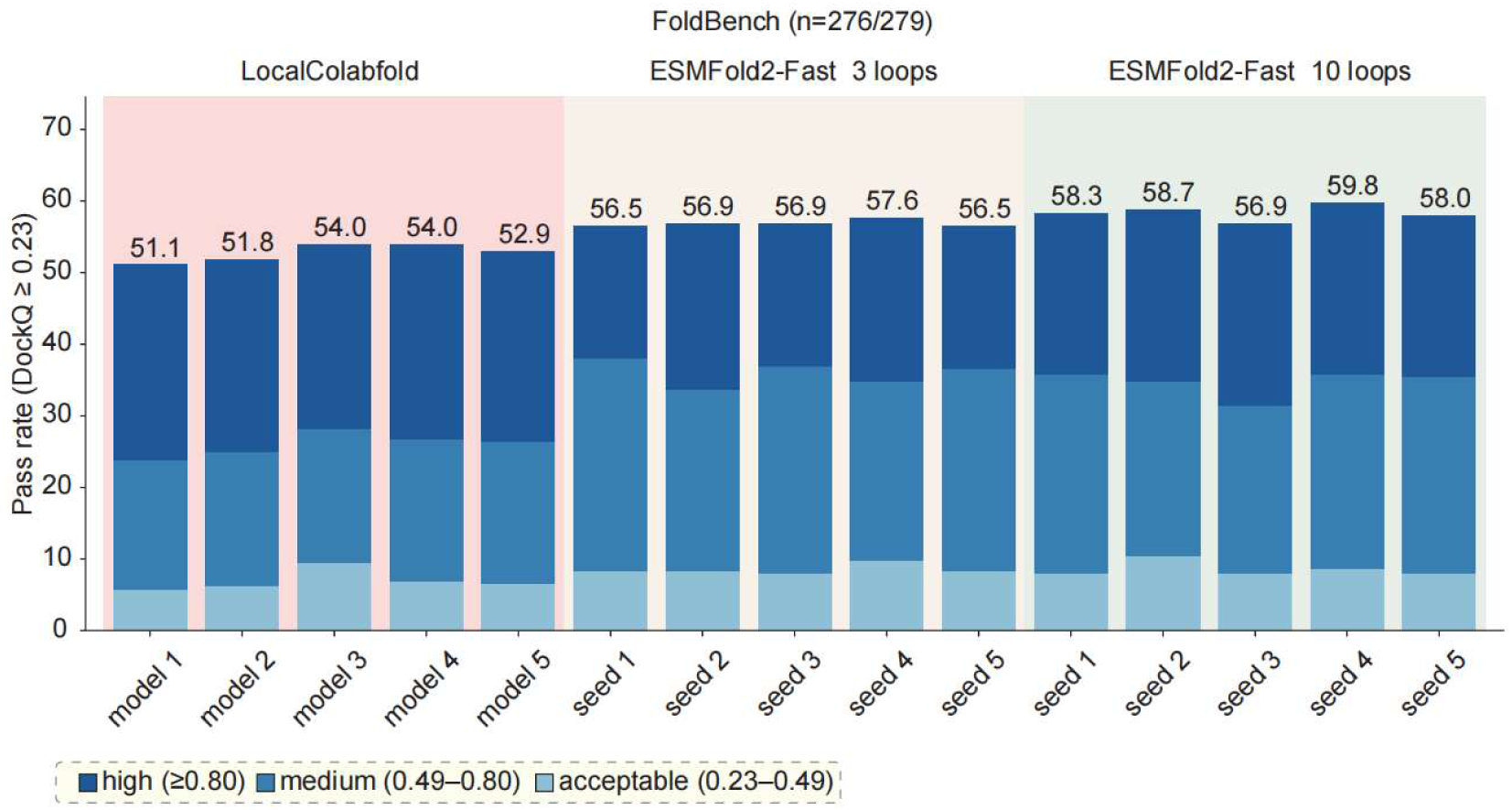
Per-output DockQ performance on FoldBench, corresponding to Figure 1A. Stacked bar plots showing the DockQ pass rate of each individual prediction output on the FoldBench protein–protein dataset (n = 276/279), for the three settings compared in Figure 1A: LocalColabFold (5 model outputs, model 1–5), ESMFold2-Fast (3 loops, seed 1–5) and ESMFold2-Fast (10 loops, seed 1–5). Each output (model or seed) was evaluated independently. For each output, the bar height gives the fraction of targets with DockQ ≥ 0.23, segmented by DockQ category: acceptable (0.23–0.49), medium (0.49–0.80) and high (≥0.80). Numbers above bars denote the overall pass rate (%).

**Figure S2.**
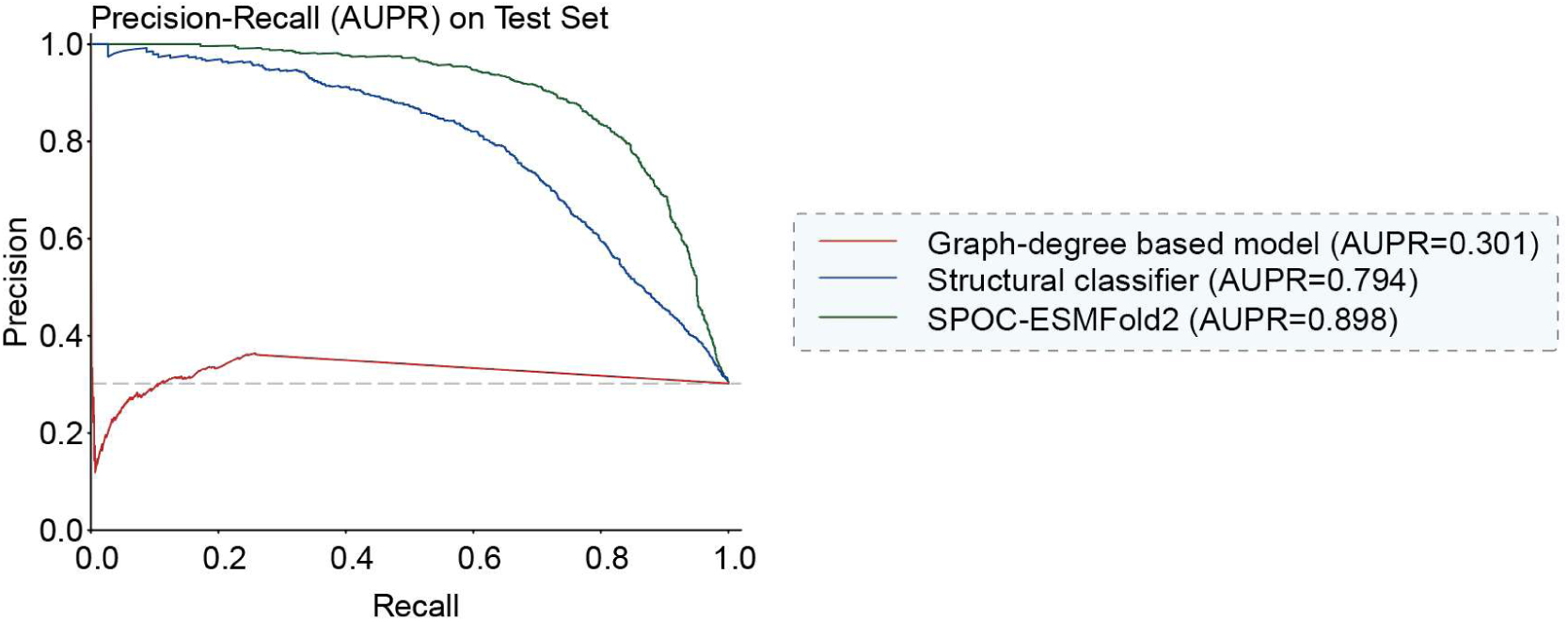
Precision–recall performance of the reranking classifiers. Precision–recall curves showing precision as a function of recall for SPOC-ESMFold2, the structural classifier, and the graph-degree based model on the held-out test set. The dashed line indicates random ranking, whose AUPR equals the positive-class prevalence.

**Figure S3.**
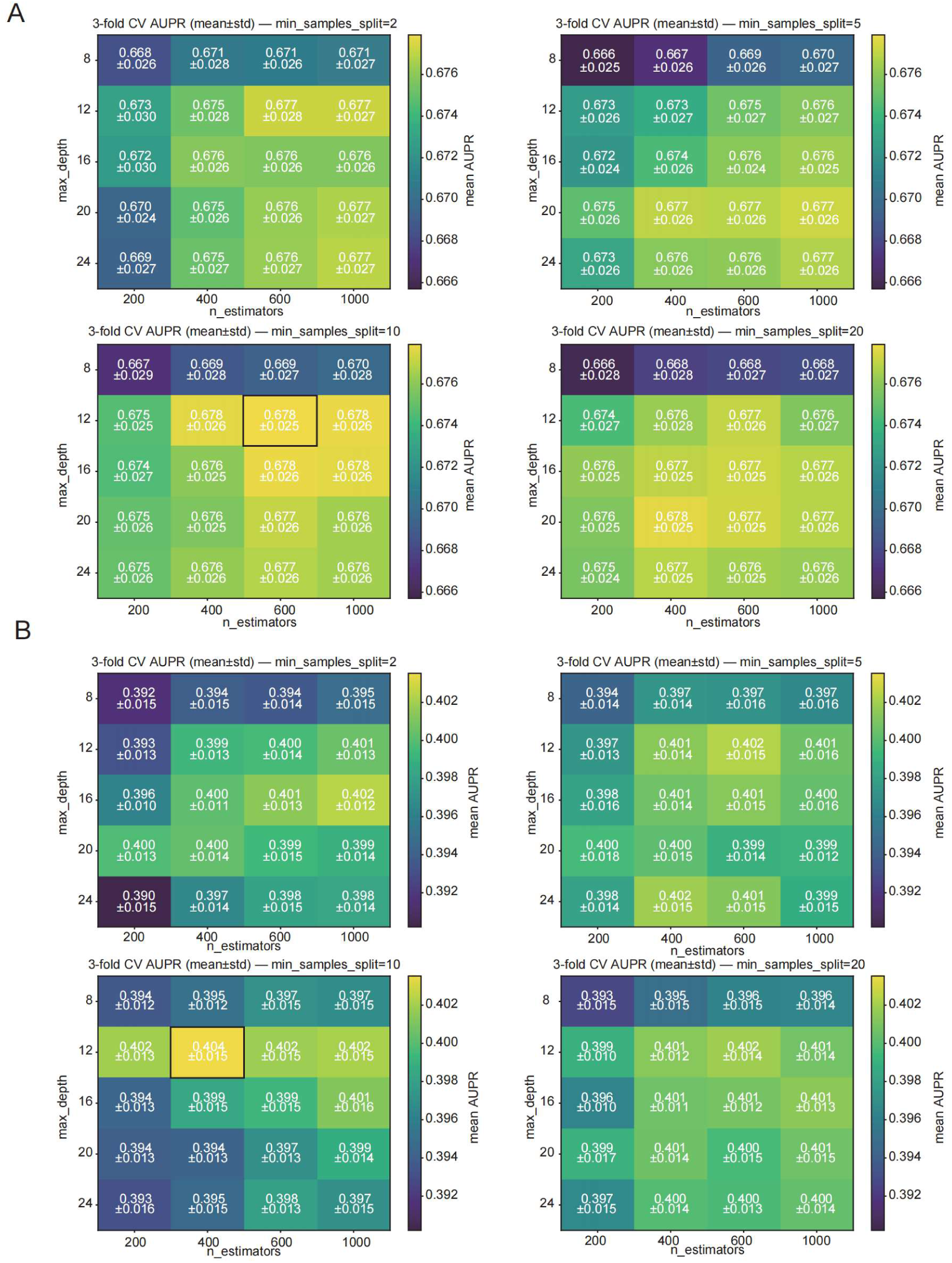
Hyperparameter tuning of the random-forest classifiers by cross-validated grid search. **(A)** Heatmaps showing 3-fold cross-validated AUPR (mean ± standard deviation) for the SPOC-ESMFold2 across the grid of n_estimators (200–1,000) and max_depth (8–24), with one heatmap per value of min_samples_split (2, 5, 10, 20). The black outline marks the selected configuration (n_estimators = 600, max_depth = 12, min_samples_split = 10; AUPR = 0.678 ± 0.025). **(B)** Same as (A) for structural classifier. The black outline marks the selected configuration (n_estimators = 400, max_depth = 12, min_samples_split = 10; AUPR = 0.404 ± 0.015).

**Figure S4.**
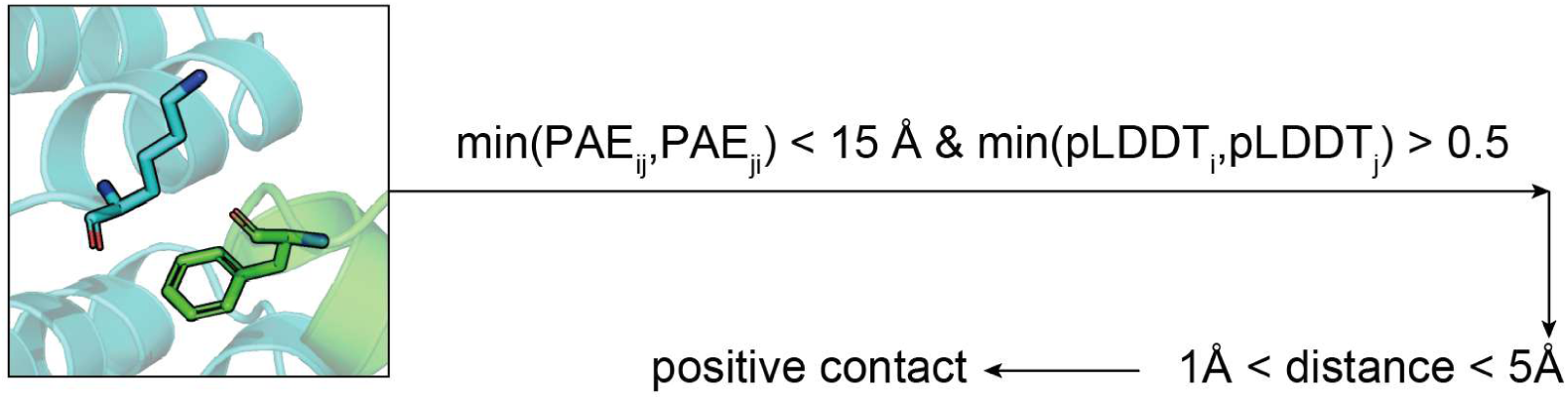
The definition of positive contacting residue. A residue pair was classified as a positive contact when three criteria were met simultaneously: PAE < 15 Å for both residues (PAEᵢ < 15 Å and PAEⱼ < 15 Å), pLDDT > 0.5 for both residues, and a closest heavy-atom distance between residue pair between 1 and 5 Å.

